# Stem cell culture conditions affect in vitro differentiation potential and efficiency of mouse gastruloid development

**DOI:** 10.1101/2024.05.09.593319

**Authors:** Marloes Blotenburg, Lianne Suurenbroek, Vivek Bhardwaj, Peter Zeller

**Author notes:** Correspondence (PZ), (MB).

## Abstract

By aggregating low numbers of mouse embryonic stem cells (mESCs) followed by exposure to Wnt activation, structures termed ‘gastruloids’ are produced that display an anteroposterior organisation of cell types derived from all three germ layers. Since current gastruloid protocols display considerable heterogeneity between experiments in terms of morphology, elongation efficiency, and cell type composition, we investigated whether an optimised mESC pluripotency state would provide more consistent results. By growing mESCs in different intervals of ESLIF and 2i medium the pluripotency state of cells was modulated, and mESC culture as well as the resulting gastruloids were analysed. Microscopy analysis showed a pre-culture-specific effect on gastruloid formation, in terms of efficiency, elongation index and reproducibility. RNA-seq analysis of the mESC start population confirmed that short-term pulses of 2i and ESLIF modulate the pluripotency state, and result in different cellular states. With multiple epigenetic regulators among the top differentially expressed genes we further analysed genome-wide DNA methylation and H3K27me3 distributions. We observed the same divergence between conditions, most dominantly displayed in the promoter regions of developmental regulators. Lastly, when we investigated the cell type composition of gastruloids grown from these different pre-cultures, we observed that mESCs subjected to 2i-ESLIF preceding aggregation generated gastruloids more consistently, including more complex mesodermal contributions as compared to the ESLIF-only control. This indicates that optimisation of the mESCs pluripotency state allows the modulation of cell differentiation during gastruloid formation.

## Introduction

The first mouse embryonic stem cell (mESC) line was established in 1981 and derived from the inner cell mass of pre-implantation blastocysts (1,2). These cells have become crucial for studying cellular differentiation in vitro. Through directed differentiation of mESCs, many different cell types can be generated, including hepatocytes, motor neurons, and T-cells (3–5). It was further found that aggregation of mESCs into 3D spheres called embryoid bodies (EBs) causes their spontaneous differentiation into structures containing cell types from all three embryonic germ layers (6). A small fraction of EBs is known to spontaneously break symmetry, start expressing the polarisation factor Brachyury (T) and acquire spatial organisation (7). This effect can be more consistently achieved by aggregating low mESC numbers (300-600) that are pulsed with a Wnt-activator, resulting in the formation of so-called ‘gastruloids’ (8–10). Gastruloids contain cell types derived from all three germ layers, and display a high level of spatial organisation, including the development of all three body axes and collinear Hox gene expression (11–15). This makes gastruloids a useful tool for studying aspects of embryogenesis in vitro.

Importantly, gastruloids are devoid of several cell types, including extra-embryonic tissues and anterior structures such as brain precursors (11,13,14). However, through the addition of signalling molecules, gastruloids can be directed to form these cell types. For instance, anterior structures can be induced through the addition of FGF and activin A and replacement of Wnt-activation with Wnt-inhibition and early stages of heart development can be modelled by including cardiogenic factors FGF, ascorbic acid, and VEGF (16,17). Alternatively, when extra-embryonic endoderm cells are added, gastruloids proceed to form visceral endoderm (18). Lastly, through embedding gastruloids in matrigel before the elongation stage, tissue structures such as somites, the neural tube, and the gut tube are more faithfully reproduced (14,19–21). Together these studies indicate that through adaptation of the gastruloid protocol, a wide variety of cell types and even tissue structures can be established.

Generally, mESCs are maintained in a medium containing serum called ESLIF, which gives rise to a heterogeneous pool of cells. These cells are comparable to E5.5 epiblast cells and are referred to as being in a naive pluripotency state (22). Alternatively, cells can also be maintained in serum-free medium containing glycogen synthase kinase 3 (GSK3) and MAP/ERK kinase (MEK) inhibitors, commonly referred to as 2i. These cells are considerably more homogeneous and revert back to an inner cell mass (ICM)-like state referred to as ground-state pluripotency (23,24). The main differences between these two states are related to their epigenome. In particular, DNA methylation and the histone modification H3K27me3, indicative of a repressive chromatin state, show variable distributions between the two conditions. 2i-grown mESCs show a genome-wide DNA methylation level of approximately 30% and a general spread of H3K27me3 across the genome (23,25,26). In contrast, in serum-grown mESCs DNA methylation covers 80% of the genome, and H3K27me3 shows more focused distributions around promoter regions (23,27–29). The differences between pluripotency states are well characterised, and appear to affect the differentiation potential of mESCs. For example, it has been reported that the pluripotency state of mESCs affects the contribution to cell types from all germ layers upon EB formation (30). More recently, it was shown that priming mESCs beyond naive pluripotency, through addition of Activin A and FGF, impairs their ability to undergo symmetry breaking and consequently blocks gastruloid formation (31). However, different laboratories use different culture media to routinely maintain mESC cultures before differentiation. Especially concerning gastruloid culture, variability of lineage contributions and recovered cell types are observed between studies (14,19,20,31). Therefore, it is possible that more subtle differences in pluripotency state of mESCs also affect differentiation, and the observed variability in differentiation assays partially results from this. Here, we applied different combinations of 2i and ESLIF culture media to mESCs, profiled their pluripotency state, and studied the effects on gastruloid differentiation and lineage contributions.

## Materials and Methods

### Cell culture and gastruloid generation

129S1/SvImJ / C57BL/6 mESCs (32) were cultured in a humidified incubator (5 % CO2, 37 °C) on 0.1 % gelatin-coated cell culture dishes in ESLIF medium (GMEM (Gibco) containing 10 % fetal bovine serum (FBS), 1 % Sodium Pyruvate, 1 % non-essential amino acids, 1 % GlutaMAX, 1 % penicillin-streptomycin, 0.2 % b-mercaptoethanol, and 1000 units/mL mouse leukaemia inhibitory factor (mLIF, ESGRO)). Cells were split every second day by washing with PBS0, dissociating with TrypLE for 5 mins at 37 °C, centrifuging at 300 g for 3 mins, and replating the pellet 1:5 on gelatinised plates. Pre-culture conditions consisted of different pulse timings and lengths with 2i medium (48.1 % DMEM/F12 (Gibco) and 48.1 % Neurobasal (Gibco) containing 0.5 % N-2 supplement (Gibco, # 17502048), 1 % B-27 supplement (Gibco, # 17504044), 1 % GlutaMAX, 1.1 % penicillin-streptomycin, 0.2 % b-mercaptoethanol, 1000 units/mL mouse leukaemia inhibitory factor (mLIF, ESGRO), 0.03 % chiron (CHIR99021, Tocris # 4423) and 0.01 % PD032509 (Sigma, # PZ0162)). During pre-culture conditions, cells were split at day 1 and 3; medium was refreshed at day 2 and 4. Gastruloids were generated as described before (11,14) with the following adaptations: N2B27 medium (Takara) was passed through a .22 µm filter before use and 600 cells were used for aggregation instead of 300.

### Bright-field microscopy and analysis

Gastruloids were kept in 96-well plates and imaged using the Leica THUNDER Imager Live Cell & 3D Cell Culture & 3D Assay with a 10x dry lens. During imaging, the temperature was kept at 37 °C and the CO2 level at 5 %. Multiple Z-stacks were taken of each gastruloid with an interval of 5 µm.

Images were analysed with the ImageJ data processing package Fiji (v.2.1.0/1.53c). Z-stacks were focused using the plug-in Stack Focuser. Cells were then segmented using the pixel segmentation function of ilastik (v.1.1.3post3), which was trained on 5 samples per condition. Next, circles were inscribed and the spine was drawn using the Max Inscribed Circles Fiji plug-in. The diameter of the largest circle and the spine length were measured with Fiji. The elongation index was calculated in Python 3.0, which was also used for further analysis. Statistics were calculated with `statannot`’s `add_stat_annot` (v0.2.3). For comparison of 2 conditions an independent T-test was used, and for more than two conditions the Mann-Whitney-Wilcoxon two-sided test with Bonferroni multiple testing correction was used.

### Dissociation, FACS sorting and scRNA-seq of mESCs and single gastruloids

mESCs were dissociated as described, re-suspended in PBS0 supplemented with 5 % FBS, and filtered through a 35 µm filter (Falcon, 352235). Single gastruloids were washed with PBS0, followed by incubation with 50 µL trypsin without phenol red (Gibco) for 5 mins at 37 °C. Gastruloids were mechanically broken up into a single-cell suspension by pipetting with a P200 pipette, diluted to 1 mL with PBS0 supplemented with 5 % FBS, and filtered through a 35 µm filter (Falcon, 352235). Prior to fluorescence-activated cell sorting (FACS), DAPI was added to the cell suspension to assess cell viability. Libraries were prepared following the CEL-seq2 protocol (33). The libraries were sequenced with the Illumina NextSeq 500 with 2 x 75 bp reads or 1 x 75 bp reads (26 bp at read 1 and 61 bp at read 2).

### sortChIC and bisulfite sequencing of mESCs

H3K27me3 and DNA methylation was assayed in bulk from the same cell suspension as used for CEL-seq2. For bisulfite sequencing, cells were pelleted after dissociation by centrifuging at 300 g for 3 mins, followed by DNA extraction using the Qiagen DNeasy Blood & Tissue Kit (69504) according to the manufacturer’s guidelines. DNA yield was quantified with Qubit dsDNA HS Assay Kit (10616763) and 100 ng DNA was used as input for the Zymo Research Pico Methyl-Seq Library Prep Kit (D5456). Abundance and quality of the final library were assessed by Qubit dsDNA HS Assay Kit (10616763) and Agilent Bioanalyzer HS DNA kit (5067-4626) and sequenced paired-end 2 x 150 bp using an Illumina NextSeq 500. For sortChIC, dissociated cells were ethanol-fixed and incubated with 1:100 anti-H3K27me3 antibody (NEB, 9733S) as described previously (34). 1000 cells were sorted in a tube and processed as described, with all single-cell volumes multiplied by 100 (34). Samples were sequenced paired-end 2 x 100 bp on an Illumina NovaSeq 6000.

### Transcriptome analysis

Raw fastq files were processed into count tables using the transcriptome Snakemake (35) pipeline of the SingleCellMultiOmics package (see https://github.com/BuysDB/SingleCellMultiOmics/wiki/Transcriptome-data-processing, version 0.1.30). The pipeline performs demultiplexing for CEL-Seq2 barcodes with a hamming distance of 0 using SingleCellMultiOmics’ demux.py and trimming with cutadapt (36) (version 4.1), followed by mapping with STAR (37) (version 2.5.3a) to the 129/B6 SNP-masked GRCm38 mouse genome (Ensembl 97). Feature counting was performed with featureCounts (38) (version 1.6.2) and reads were tagged and deduplicated, then transformed to a .csv file. Downstream analysis was performed with Scanpy (39) (version 1.8.1). Cells with either less than 500 reads and 100 genes detected in less than 2 cells were filtered out. Cells with more than 40 % or more than 20 % mitochondrial reads were excluded for mESCs and gastruloid cells, respectively. Mitochondrial reads, Malat1, and ribosomal reads were excluded from further analysis. Counts were normalised to 500 transcripts per cell and logarithmized. The number of counts, percentage of mitochondrial reads, and cell cycle phase were regressed out and each gene was scaled to unit variance with a maximum of 10. Principal component analysis was performed with 10 neighbours and either the 30 or 40 highest principal components were used to generate the UMAP for mESCs or gastruloid cells, respectively. The data was clustered using the Leiden algorithm (scanpy.tl.leiden, resolution set to 0.5). Differentially expressed genes between clusters were determined using Scanpy’s Wilcoxon rank-sum test. Differentially expressed genes between ESLIF and 2i pre-culture states were determined by grouping pre-culture conditions into pseudobulks with decoupler (version 1.6.0) followed by pydeseq2 (version 0.4.4)

### H3K27me3 and DNA methylation data analysis

WGBS-seq data was mapped using the snakePipes (40) WGBS mapping workflow (v 1.3.1) to GRCm38 (mm10) genome with default parameters. WGBS workflow applies paired-end trimming using fastp (41) (v0.20) with options -q 5 -l 30 -M 5, mapping using bwa mem (42) with options -T 40 -B 2 -L 10 -CM -U 100 -p and bwa-meth (43) (v0.2.2) for methylation encoding, followed by CpG methylation calling using methyldackyl (https://github.com/dpryan79/MethylDackel, v0.5.0) with options -mergeContext -maxVariantFrac 0.25 -minDepth 4. ChIC-seq data was mapped using the ChIC Snakemake pipeline (35) of the SingleCellMultiOmics package (version 0.1.30). The CPM-normalised coverage was computed in 100 bp bins using deepTools (44) (v3.0) bamCoverage with options -normalizeUsing CPM -skipNAs -ignore X MT. We used the mm10 gencode (45) (v23) annotation to define promoter (TSS +-10 kb), gene body (signal >10 kb downstream of TSS, only on genes longer than 10 kb), intergenic regions and repbase database (2022) to define repeat regions. WGBS signal (CpG methylation ratio) and ChIC signal (counts) were then aggregated on these grouped features. In order to compare ChIC signal between features, we additionally normalised the ChIC signal to the total length of the feature groups using FPM (features per million) defined as FPM = A * (1 / (sum(A))) * 10 x 106, where A = feature counts/kb. Finally, to classify genes based on the ChIC signal at the promoters, we aggregated counts on promoter regions using computeMatrix with options -bs 200 –missingDataAsZero -skipZeros, followed by plotHeatmap with options -kmeans 4 to get the 4 gene clusters. The normalised counts per cluster was then calculated as CPM = A / (sum(A) * 10 x 106), where A = feature counts.

### Code and data availability

All data generated for this project, code used and annotated notebooks for performing the analysis and generating figures can be found on github (https://github.com/marloes3105/preculture).

## Results

### Different intervals of 2i exposure during mESC culture affect the reproducibility of gastruloid formation

To test the impact of preculture-defined mESC states on gastruloid composition, we applied different combinations of 2i and ESLIF culture media to mESCs and studied the effects on gastruloid formation (Fig 1A). For this study, we used an mESC line with a 129SV/Bl6 background, which we routinely maintained in ESLIF medium on gelatin (32). We included two control conditions: one with ESLIF only (1) and one with 2i only (2), representing naive and ground state pluripotency, resp., and show the expected morphologies, with flat and connected colonies in ESLIF medium and dome-shaped disconnected colonies in 2i medium (Fig 1A’). We hypothesised that pluripotency states in between naive and ground state can be produced with short-term pulses of 2i and ESLIF. To this end, we tested 4 intermediate states. The morphologies of mESCs subjected to the different pulses are in between the ESLIF and 2i state, with conditions 3 and 5 forming ESLIF-type colonies with round edges, condition 4 resembling the ESLIF control, and condition 6 forming large dome-shaped colonies (Fig 1A’).

**Figure 1.**
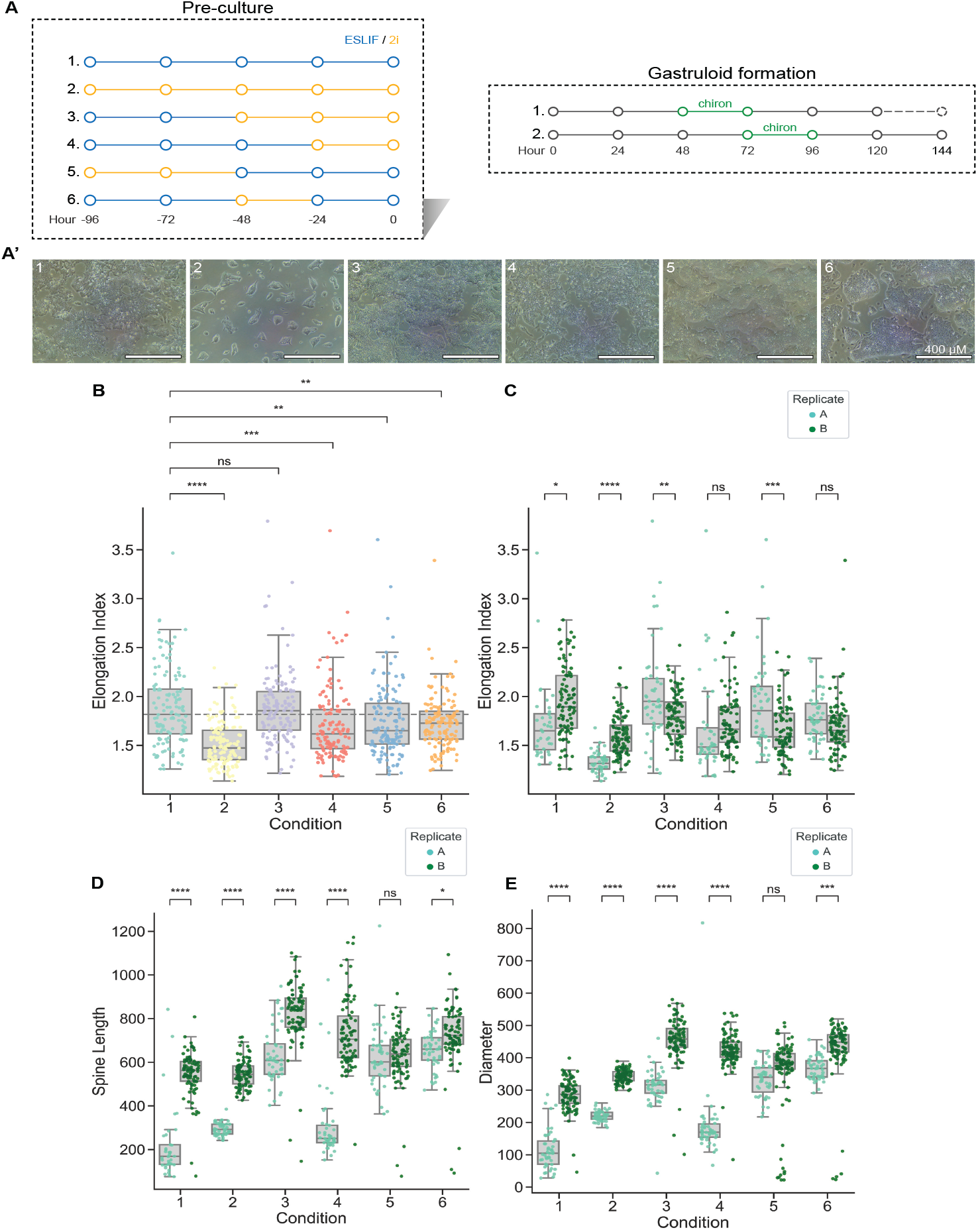
Variations in mESC culture conditions affect gastruloid elongation length, efficiency, and consistency across replicates. (**A**) Schematic overview of six different pre-culture conditions, with varying times of ESLIF or 2i culture medium, and (A’) corresponding bright-field images showing cell morphologies for all conditions. Gastruloid formation was performed according to the standard protocol (1) or using a 24-hour delayed chiron pulse and read out time point (2). (**B**) Elongation index of gastruloids. The most efficient timing of chiron pulse and read-out was chosen for each condition (see Figs S1C-E). For condition 3, this was 24 h delayed (chiron pulse at 72-96 h AA, read-out at 144 h AA), for all other conditions, this was the standard protocol (chiron pulse at 48-72 h AA, read-out at 120 h AA). (**C**-**E**) Elongation index (**C**), spine length (**D**) and diameter (**E**) of produced gastruloids, separated by replicate. Statistical significance between replicates A and B was calculated with an independent t-test.

mESCs were aggregated into gastruloids following the standard protocol which uses a Wnt-activation pulse between 48 h and 72 h after aggregation (AA) through the addition of chiron (11,14). Gastruloids display morphological variation at 120h AA depending on the mESC pre-culture condition used (Fig S1A). Notably, cells from all pre-culture conditions except condition 2 generate gastruloids efficiently. Cells cultured in condition 3 and 5 develop into gastruloids with more complex internal structures, such as a ridge in the centre and protrusions at the posterior end (Fig S1A). Because some pre-culture conditions could be delayed in becoming perceptive to the chiron pulse due to their pluripotency state, we subjected all conditions additionally to a delayed chiron pulse from 72 h - 96 h AA, and assessed them at 144 h AA (Fig 1A). To characterise a high number of gastruloids in parallel, we developed a gastruloid elongation screening. In short, we imaged all gastruloids generated, and developed an ImageJ macro to extract the spine length and diameter of each individual gastruloid to calculate the elongation index (Fig S1B). This approach was used in all following analyses.

While gastruloids from most pre-culture conditions show a lower elongation index following the delayed chiron pulse, condition 3 cells display a significant increase in elongation after a delayed pulse and delayed read-out (Figs S1C-E). Therefore, we subjected condition 3 pre-culture cells to a 72 h - 96 h AA chiron pulse and read-out at 144 h AA, while for all other pre-culture mESCs we used the default gastruloid protocol.

Only gastruloids generated from condition 3 mESCs did not produce a significantly lower index than the ESLIF-only control (condition 1), while 2i-state mESCs (condition 2) result in gastruloids with the lowest elongation index (Fig 1B). When we split the data by experiment, we observed a high degree of heterogeneity between replicates except for conditions 4 and 6 (Fig 1C). When looking at the absolute length and diameter of the gastruloids, this effect is further enlarged (Figs 1D-E). Only gastruloids generated from pre-culture conditions 5 and 6 show consistent results across the two replicates, and only condition 5 does not show significant differences in any of the measured parameters (Figs 1D-E).

To conclude, the pre-culture conditions to which mESCs are exposed indeed affect the formation of gastruloids. Variations in mESC pre-culture result in gastruloids with different morphologies and elongation scores (Fig 1). Finally, pre-culture conditions of mESCs affect reproducibility of gastruloid generation, especially when observing the spine length and diameter scores, with only pre-cultures 5 and 6 generating reproducible results (Figs 1D-E).

### mESC pre-culture conditions display two distinct transcriptional and epigenetic signatures

Next, we wondered if the observed phenotypic variability during gastrulation is already reflected in the mESC state during aggregation. Single-cell RNA sequencing analysis of the mESCs showed a clear segregation between cells from conditions 1, 5, and 6 on the one hand, and pre-culture conditions 2, 3, and 4 on the other (Fig 2A). This effect is not due to plate effects (Fig S2A) or other technical aspects, such as the total genes profiled, total counts recovered, and percentage of mitochondrial reads per cell (Figs S2B-D). In contrast to the observed variability in gastruloid formation, mESCs inside each condition were relatively homogenous. Next, we clustered the cells into cell states and determined the top differentially expressed genes for each cluster using the Wilcoxon rank-sum test. Cell state clusters are largely dominated by cells from the same pre-culture condition, with the exception of two outlier clusters (Fig 2B). These two clusters also show a lower S-phase score compared to the other clusters, indicating a low cell division rate (Fig S2E). We classified one of the two outlier clusters as ‘Endoderm Progenitors’, due to high and specific expression of endoderm-related genes including Foxa2, Sox17, Sox7, Gata4, and Gata6 (Fig S2F). The second outlier cluster shows expression of both mesoderm specification (Mesp2, Notch1) and cell cycle arrest (Ccng1, p53), consistent with a lower S-phase score, indicative of Mesoderm Progenitors in cell cycle arrest (Figs S2E-F).

**Figure 2.**
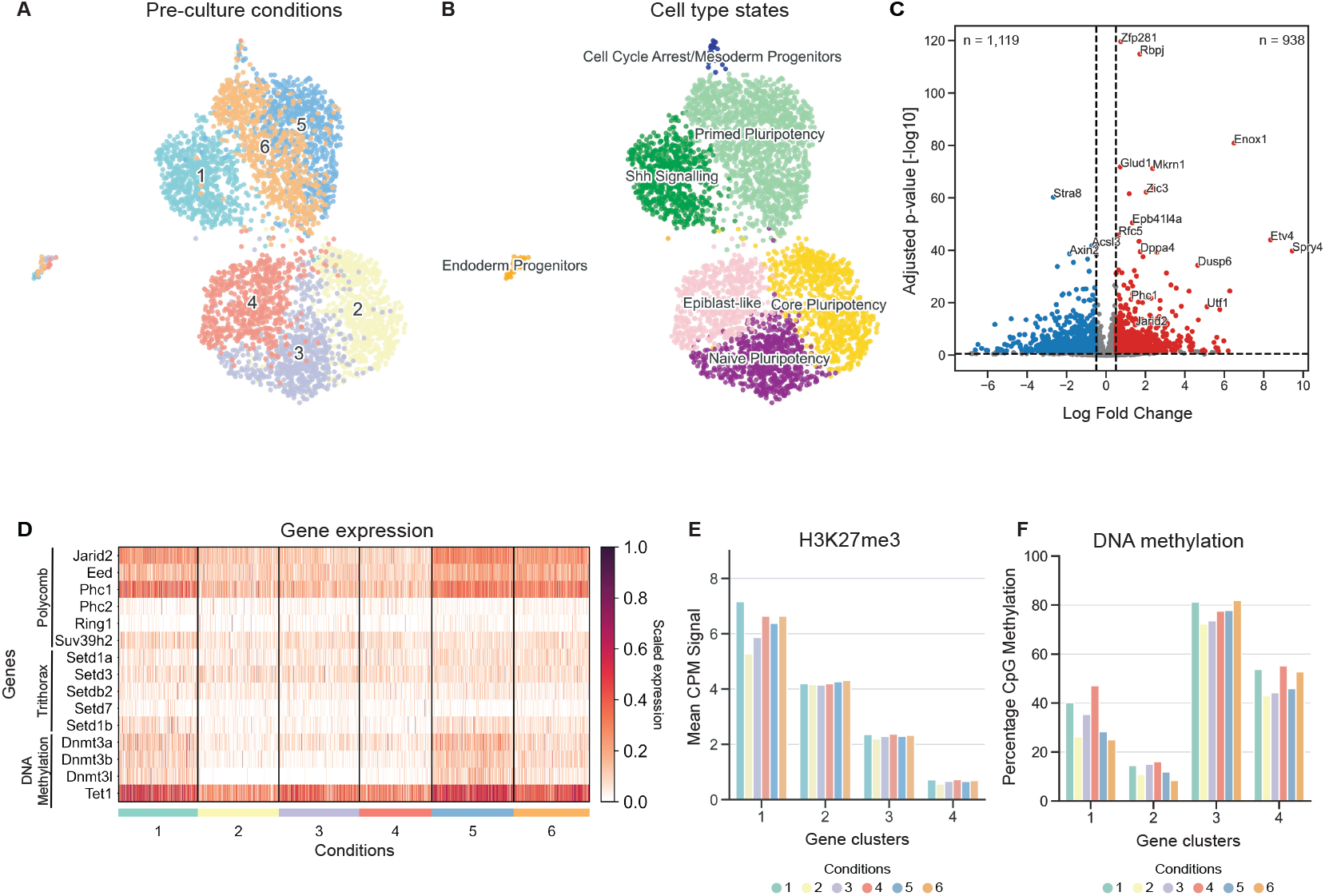
Specific pulses of mESC culture media result in distinct transcriptome profiles and differential gene expression of epigenetic marker genes. (**A**) UMAP clustering of single-cell transcriptomes (n = 4,147) of mESCs cultured using pre-culture conditions defined in Fig1. (**B**) UMAP coloured by cell type states: Primed Pluripotency (n = 1,396), Shh Signalling indicative of ectodermal priming (n = 658), Mesoderm Progenitors/cells in Cell Cycle Arrest (n = 28), Endoderm Progenitors (n = 45), Epiblast-like cells (n = 602), cells in Core Pluripotency state (n = 771), and cells in Naive Pluripotency state (n = 647). Cell typing is based on marker gene expression displayed in Fig S2F. (**C**) Volcano plot indicating downregulated genes (left, blue) and upregulated genes (right, red) in ESLIF pre-culture conditions (conditions 1, 5, 6) compared to 2i pre-culture conditions (conditions 2, 3, 4). (**D**) Heatmap showing differential expression between the pre-culture conditions of Polycomb-associated genes (Jarid2, Eed, Phc1, Phc2, Ring1, Suv39h2), DNA methylation factors (Dnmt3a, Dnmt3b, Dnmt3l, Tet1), and Trithorax-associated genes (Setd1a, Setd3, Setdb2, Setd7, Setd1b). (**E**) H3K27me3 abundance in a 20 kb window around the TSS of genes grouped by differential gene clusters. Differential gene clusters were identified based on their H3K27me3 TSS coverage across the six conditions (**F**) Percentage of DNA methylation on promoter CpGs, grouped per pre-culture condition and gene cluster determined based on H3K27me3 data.

The majority of the cells display high expression of pluripotency genes including Sox2, Esrrb, and Nanog (Fig S2F). However, we could discern differences between pluripotency states. 2i-grown mESCs (condition 2) show high expression of Core Pluripotency genes Nanog, Epha4, and Tex19.1, while condition 3 cells exhibit a Naive Pluripotency signature with Gjb3, Hes1, and Klf4, and condition 4 mESCs resemble Epiblast-like cells with up-regulation of Igfbp2, Tuba1a, and Fos (Fig S2F). mESCs grown in pre-culture conditions 5 and 6 show a Primed Pluripotency signature with expression of Nr0b1, Tet2, Dppa2, Dppa4, and Bmp4, and condition 1 mESCs show an ectodermal signature with expression of Gbx2, Ptch1, Sox11, and Lefty1, indicative of Sonic Hedgehog (Shh) signalling (Fig S2F). Calculated correlation scores confirmed the split between pre-culture conditions (Fig S2G) and confirmed Endoderm Progenitors and Cell Cycle Arrest/Mesoderm Progenitors as outlier populations that likely represent a small fraction of differentiated cells (Figs 2B, S2H). To look into this split between the pre-culture conditions, we performed differential gene analysis between conditions 1, 5, and 6 (ESLIF conditions), compared to 2, 3, and 4 (2i conditions). Top differentially expressed genes include known pluripotency factors as well as epigenetic factors such as Utf1, Jarid2, and Phc1 (Fig 2C). Epigenetic changes between 2i-grown and ESLIF-grown mESCs are well studied, in terms of DNA methylation and chromatin modifications (22,23,25–29). While H3K27me3 is deposited by the Polycomb Repressive Complex (PRC), H3K4me3 is regulated through Trithorax Group Proteins (TxG). Indeed, when we analysed gene expression patterns of Polycomb, Trithorax and DNA methylation factors, we observed an up-regulation of PRC factors in conditions 1, 5, and 6, but not in 2, 3, and 4 (Fig 2D). This same pattern is to a lesser extent replicated in DNA methylation factors, but not in TxG factors (Fig 2D).

To look further into epigenetic differences between the conditions, we profiled DNA methylation with Whole Genome Bisulfite Sequencing (WGBS) and H3K27me3 distribution across the genome with Sort-Assisted Chromatin Immunocleavage (sortChIC) (34). First, we assessed distribution of both modifications across different genomic regions. While repeat regions are generally highly covered by H3K27me3, gene bodies and promoters contain a lower amount of this mark (Fig S2I). DNA methylation is generally highest in intergenic regions and lowest in promoter regions (Fig S2J). We observed the largest variation between pre-culture conditions in promoter regions, and therefore focused the following analyses on these (Fig S2J). We assessed coverage of H3K27me3 in a 20kb window around individual transcription start sites (TSS) and identified four gene clusters with variable H3K27me3 distribution dynamics (Fig 2E). To assess the effects of these epigenetic differences on gene expression, we grouped the scRNA-seq dataset generated before according to the same gene clusters and analysed the average expression per pre-culture condition and gene cluster (Fig S2K). Around 20% of the genes classified to clusters 1 and 2 are differentially expressed between the ESLIF and 2i conditions, with more genes being downregulated (14.5% and 12.4%, resp.) than upregulated (8.3% and 9.4%, resp., Fig S2L).

Generally, gene cluster 1 shows variability in the amount of DNA methylation and H3K27me3 coverage between pre-culture conditions, but not when assessing gene expression (Figs 2E-F, S2K). GO analysis reveals that the variable gene cluster 1 is associated with terms like system development, multicellular organism development, and cell differentiation (Fig S2M). In contrast, while genes classified in cluster 2 show generally high levels of H3K27me3 coverage, DNA methylation across these gene promoters is low with levels below 20% (Figs 2E-F). Genes classified to gene clusters 1 and 2 are generally lowly expressed, consistent with their silencing through H3K27me3 (Figs 2E-F, S2K). Genes classified to cluster 3 follow the inverse pattern with DNA methylation levels reaching 80%, intermediate levels of expression and corresponding lack of H3K27me3. Lastly, genes classified to cluster 4 show lower levels of both DNA methylation and H3K27me3 coverage, consistent with generally high levels of gene expression (Figs 2E-F, S2K).

### mESC pre-culture conditions affect lineage contribution in gastruloids

To study the effect of different mESC pre-culture conditions on the differentiation potential, we generated gastruloids from pre-culture conditions 1, 5, and 6 as described above. Pre-culture condition 1 was selected as a control, and conditions 5 and 6 were selected because they gave the most consistent results with high gastruloid formation efficiency in previous assays (Figs 1-2, S1-S2). Considering their similar mESC profile and gastruloid morphologies upon differentiation, we hypothesised that conditions 5 and 6 would also yield comparable cell type compositions, while condition 1 might show a bias towards ectoderm consistent with its priming through Shh signalling in mESCs (Figs S1A, 2B, S2F) . For each pre-culture condition, we selected 5 single gastruloids and subjected them to single-cell RNA sequencing to evaluate reproducibility. After excluding low-quality cells (see Methods), the remaining 4,637 single cells were used for UMAP embedding (Figs 3A, S3A). All 3 conditions contain cell types derived from all three germ layers, with mesoderm contributing the highest and endoderm contributing the lowest number of cells (Figs 3B-C, S3B). Quality metrics including the total number of genes profiled, total counts recovered, and percentage of mitochondrial reads per cell are comparable between conditions and cell types (Fig S3C).

**Figure 3.**
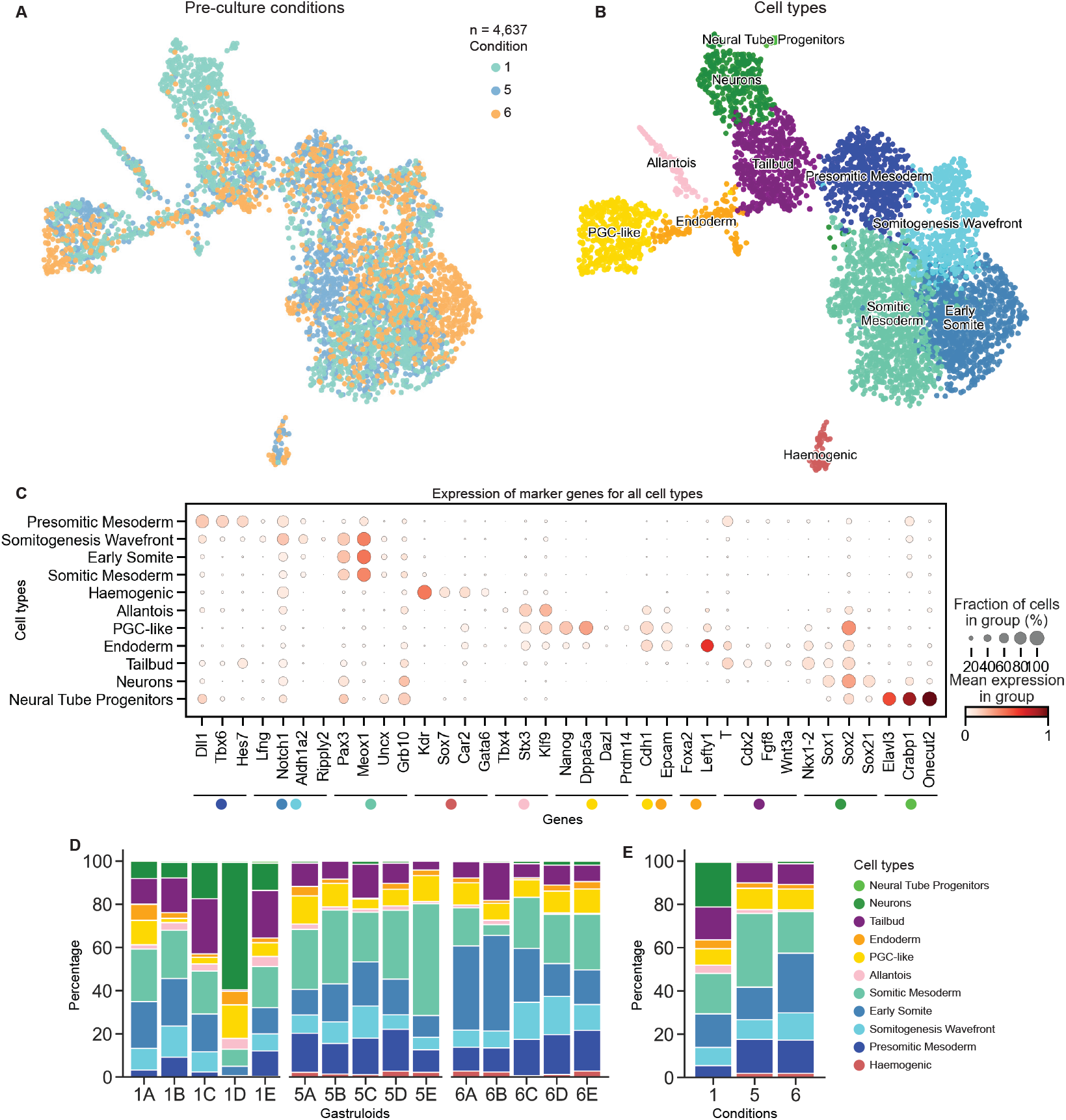
mESC pre-culture conditions affect lineage contribution in gastruloids. (**A**) UMAP showing clustering of single-cell RNA sequencing data of five individual gastruloids for conditions 1 (n = 1,529), 5 (n = 1,532), and 6 (n = 1,576), sampling a total of 4,637 cells. (**B**) UMAP coloured by recovered cell types: Primordial Germ Cell-like Cells (PGC-like, n = 419); Allantois cells (n = 96); Endodermal cells (n = 134); mesodermal cell types, including Early Somites (n = 903), Somitic Mesoderm (n = 1,111), Presomitic Mesoderm (n = 563), Somitogenesis Wavefront (n = 467), Haemogenic cells (n = 62), Tailbud cells (n = 529); and ectodermal cell types including Neurons (n = 344), and Neural Tube Progenitors (n = 9). Cell type annotation is based on localised expression of marker genes (see Fig C). (**C**) Dot plot showing the expression of marker genes in all cell types. Dot colour represents the mean expression of marker genes in specific groups, and dot size represents the fraction of cells within the group that show expression of the gene. (**D**) Cell type distributions across 5 individually sampled gastruloids from condition 1 (1A-E), condition 5 (5A-E), and condition 6 (6A-E). (**E**) Average cell type distributions from (**D**) between conditions 1, 5, and 6.

When we assessed the ratio of cell types for each gastruloid derived from each condition, we observed that condition 1 is skewed towards an ectodermal fate and generates a significantly higher ratio of Neurons (Figs 3D-E, S3D-F). This is consistent with a bias towards ectoderm in condition 1 mESCs, identified by upregulation of Shh signalling factors (Figs 2B, S2F). In contrast, gastruloids derived from conditions 5 and 6 generate a significantly higher fraction of mesoderm cells, with a bias towards Somitic Mesoderm and Early Somites, respectively (Figs 3D-E, S3D-F). Lastly, while condition 1 gastruloids produce the rare ectodermal cell type Neural Tube Progenitors, condition 5 and 6 gastruloids generate Haemogenic Progenitors (Figs S3E-F).

## Discussion

Here, we have shown that by modulating the pluripotency state of mESCs through pre-culture media composition, the efficiency and reproducibility of gastruloid formation is affected. While in our study 2i-grown mESCs yielded the lowest efficiency of gastruloid elongation, another study using E14 mESCs reported increased efficiency of elongation after mESCs were maintained in 2i medium (31). It is therefore plausible that different cell lines require alternative pre-treatments to achieve the same pluripotency state and thereby a higher efficiency upon differentiation. In addition, we observe that variations in pre-culture conditions could lead to a delayed perceptiveness to the Wnt activation pulse upon gastruloid formation, and propose to take this into consideration when performing the gastruloid protocol with untested cell lines.

Next, we suggest that modulating the pluripotency state can also affect downstream lineage potential upon differentiation, consistent with current literature (30). Concerning gastruloid differentiation, a study reporting mESC maintenance in ESLIF medium identified a high ratio of ectoderm cells, while another study routinely maintained mESCs in 2i-medium and reported faithful emergence of endoderm cells (14,20). In our study, synchronising mESCs with a 2i-pulse before exposing them to ESLIF followed by aggregation yielded a higher fraction of mesoderm compared to the ESLIF-only control. Maintenance of mESCs in ESLIF produced a higher ratio of ectodermal cells in gastruloids, consistent with ectodermal priming through Shh signalling observed under these conditions. This indicates that a higher ratio of ectoderm cells upon differentiation is already visible by early activation of Shh signalling in mESCs, and can be identified by targeted screening for the differentially expressed genes. However, overall differences between mESCs in the pre-culture conditions tested are low, both in their transcriptome and epigenetic profiles. Generally, mESC pre-cultures that generated more consistent gastruloids were marked by upregulation of epigenetic regulators of DNA methylation and H3K27me3, corresponding with differential coverage of these marks across promoter regions of developmental regulators. However, this signature did not explain the difference in cell fate composition between pre-culture 1 and pre-cultures 5 and 6. Future work looking at multiple states along differentiation might show clearer and more predictive differences.

Lastly, we present an unbiased screening approach to assess the efficiency of gastruloid formation. Through imaging of whole gastruloid plates and the quantification of morphological properties including diameter, spine length and elongation index, the effect of different treatments can be easily assessed. Using this approach, we envision that gastruloid culture can be systematically optimised for reproducibility in a wide variety of cell lines and genetic backgrounds, before a more detailed analysis of the resulting cell composition is assessed.

## Acknowledgements

The authors thank A. van Oudenaarden for supporting this study. We thank V. van Batenburg for help with the imaging experiments and for useful discussions. J. Verity-Legg assisted with bisulfite sequencing experiments. We are grateful to S. van den Brink and S. de Vries for sharing their cell culture expertise, and to D. Turner and T. van Boxtel for providing useful information and discussions. We thank the Hubrecht Sorting Facility (HSP) and Utrecht Sequencing Facility (USEQ) for cell sorting and sequencing. The 129S1/SvImJ / C57BL/6 mESC line was a gift from Matyas Flemr and Marc Bühler. P.Z. was funded by SNF (P2BSP3-174991), HFSP (LT000209/2018-L) and Marie Skłodowska-Curie Actions (798573). V.B. was funded by EMBL LTF (ALTF 1197–2019). The Oncode Institute is partially funded by the KWF Dutch Cancer Society. This work is part of the Oncode Institute which is partly financed by the KWF Dutch Cancer Society.

## Author contributions

MB designed and led the project. LS performed imaging, LS and MB performed cell culture and RNA-seq experiments, MB and PZ performed sortChIC experiments and MB performed bisulfite experiment. MB, LS and VB performed data analysis. MB and PZ wrote the manuscript with input from LS and VB. PZ supervised the study.

## Supplemental Figures

**Figure S1.**
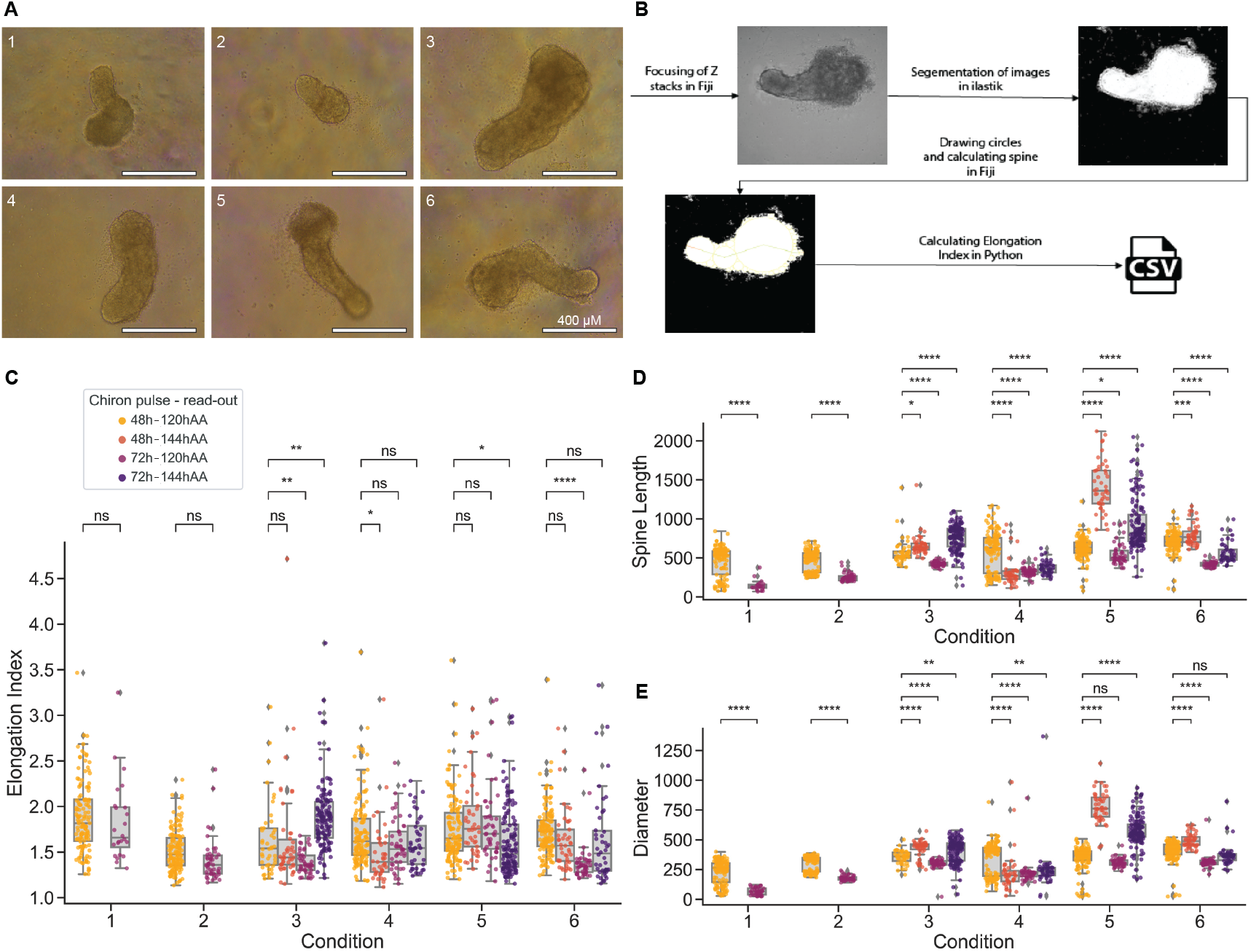
Variations in mESC pluripotency state affects gastruloid morphology, perceptiveness to chiron pulse, elongation length and efficiency of symmetry breaking. (**A**) Examples of gastruloid morphologies generated from the six different pre-culture conditions. (**B**) Workflow for quantification of spine length, diameter and elongation index of gastruloids. 96-well plates with gastruloids are imaged, and images of gastruloids are loaded into Fiji. Images were segmented using Ilastik, and circles were drawn within the shape of the gastruloids. The diameter of the largest circle was defined as the diameter of the gastruloid. A line was drawn through the circles from anterior to posterior, and the length of this line was defined as the gastruloid spine length. The elongation index was calculated by dividing the spine length by the diameter of each gastruloid. (**C**-**E**) For each pre-culture condition, the effect of conventional timing of the chiron pulse (48-72h AA: light orange, dark orange) and a delayed chiron pulse (72-96h AA: light purple, dark purple) on gastruloid formation were assessed. Gastruloid formation was assessed at 120 h AA and 144 h and the elongation index (**C**), spine length (**D**) and diameter (**E**) were calculated. Statistical significance of the differences in chiron pulse and read-out within conditions were calculated with Mann-Whitney U test with Bonferroni multiple testing correction.

**Figure S2.**
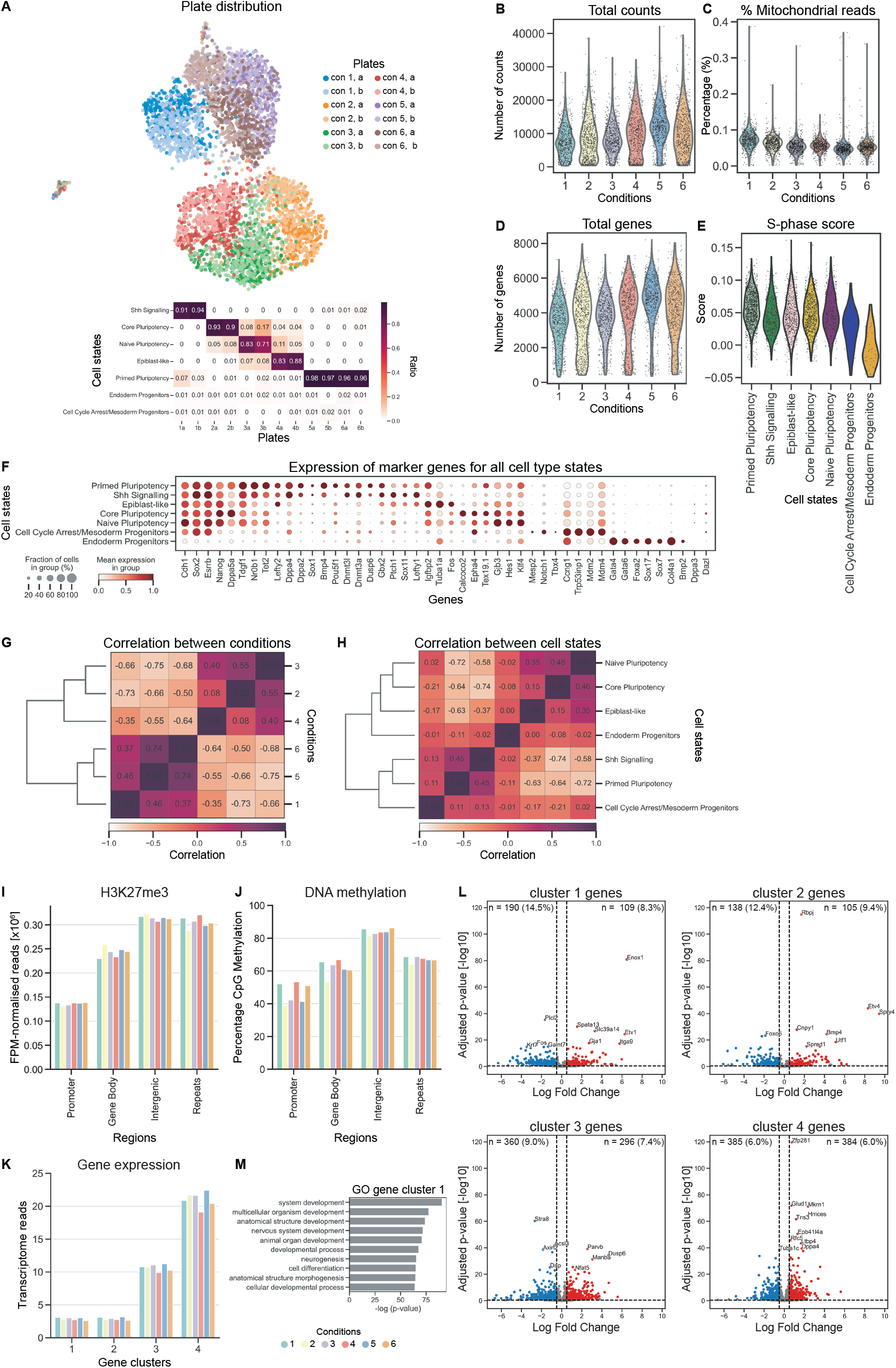
Specific pulses of mESC culture media result in distinct transcriptome and epigenetic profiles. (**A**) UMAP coloured by plate distribution, insert: ratios of cell type states per plate. (**B**)-**D**) Distribution of total count distribution (**B**), mitochondrial gene expression (**C**), and total gene count distribution (**D**) per cell. (**E**) S-phase scores per cell type. (**F**) Dot plot showing expression of marker genes used for determination of cell type states displayed in Fig 2B. Dot colour represents the mean expression of marker genes in specific groups, and dot size represents the fraction of cells within the group that show expression of the gene. (**G**-**H**) Correlation maps between conditions (**G**) and between cell type states (**H**). (**I**) Normalised H3K27me3 abundance across different genomic regions for all six mESC pre-culture conditions tested. Counts are normalised for features per million (FPM), which takes into account the variable sequencing depth across samples and the variation in genome size of the different features. (**J**) Percentage of CpG methylation across different genomic regions for all six mESC pre-culture conditions tested. (**K**) Average level of transcriptome reads derived from scRNAseq dataset, grouped per pre-culture condition and gene cluster determined based on H3K27me3 data. (**L**) Volcano plots corresponding to Fig2C separated by identified gene clusters, indicating down regulated genes (left, blue) and upregulated genes (right, red) in ESLIF pre-culture conditions (conditions 1, 5, 6) compared to 2i pre-culture conditions (conditions 2, 3, 4). (**M**) Top 10 GO terms associated with gene cluster 1.

**Figure S3.**
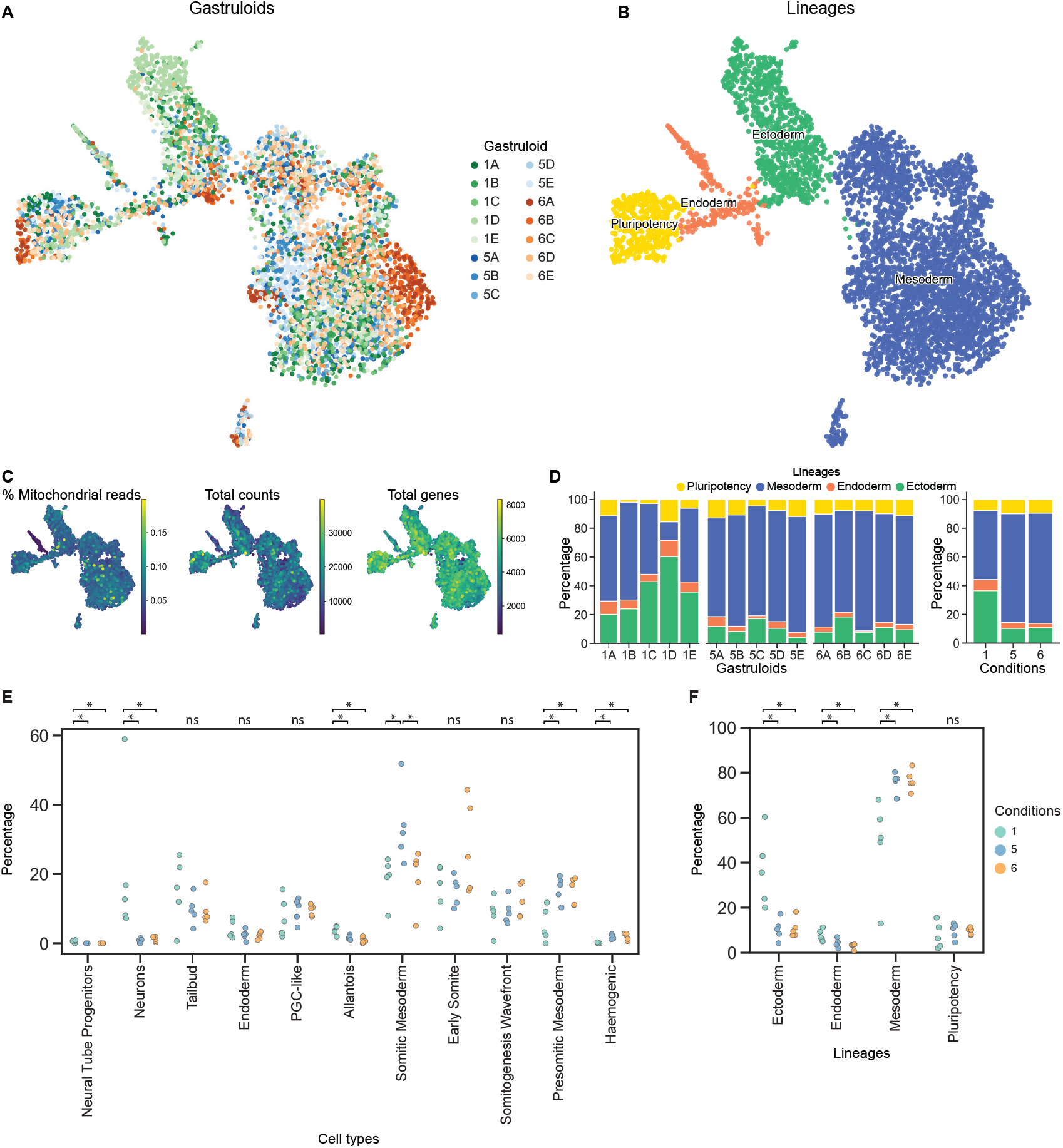
Variations in mESC pluripotency state affects cell type composition of gastruloids. (**A**) Distribution of replicates across transcriptome UMAP. Five individual gastruloids were included for pre-culture conditions 1, 5, and 6. (**B**) Transcriptome UMAP separated by germ layers - endoderm (n = 230), mesoderm (n = 3,106), ectoderm (n = 882) - and pluripotency cells (n = 419). (**C**) Distribution of mitochondrial gene expression (left), total count distribution (middle), and total gene count distribution (right) per cell. (**D**) Ratio of germ layer contributions across 15 individually sampled gastruloids from condition 1 (1A-E), condition 5 (5A-E), and condition 6 (6A-E) (left) and between conditions 1, 5, and 6 (right). (**E**-**F**) Boxplots of cell type (**E**) and lineage (**F**) percentages per condition. Statistical significance between conditions was calculated with Wilcoxon rank-sum test with Bonferroni multiple testing correction.

